# First Characterization of a Biphasic, Switch-like DNA Amplification

**DOI:** 10.1101/169805

**Authors:** Burcu Özay, Cara M Robertus, Jackson L Negri, Stephanie E McCalla

## Abstract

We report the first DNA amplification chemistry with switch-like characteristics: the chemistry is biphasic, with an expected initial phase followed by an unprecedented high gain burst of product oligonucleotide in a second phase. The first and second phases are separated by a temporary plateau, with the second phase producing 10 to 100 times more product than the first. The reaction is initiated when an oligonucleotide binds and opens a palindromic looped DNA template with two binding domains. Upon loop opening, the oligonucleotide trigger is rapidly amplified through cyclic extension and nicking of the bound trigger. Loop opening and DNA association drive the amplification reaction, such that reaction acceleration in the second phase is correlated with DNA association thermodynamics. Without a palindromic sequence, the chemistry resembles the exponential amplification reaction (EXPAR). EXPAR terminates at the initial plateau, revealing a previously unknown phenomenon that causes early reaction cessation in this popular oligonucleotide amplification reaction. Here we present two distinct types of this biphasic reaction chemistry and propose dominant reaction pathways for each type based on thermodynamic arguments. These reactions create an endogenous switch-like output that reacts to approximately 1pM oligonucleotide trigger. The chemistry is isothermal and can be adapted to respond to a broad range of input target molecules such as proteins, genomic bacterial DNA, viral DNA, and microRNA. This rapid DNA amplification reaction could potentially impact a variety of disciplines such as synthetic biology, biosensors, DNA computing, and clinical diagnostics.

## INTRODUCTION

Isothermal oligonucleotide amplification chemistries have become increasingly popular due to their simplicity and adaptability to a variety of systems^1^. Enzyme-free strand displacement amplification cascades can rapidly produce free oligonucleotides with nanomolar input trigger concentrations^2,3^. Other oligonucleotide amplification reactions rely on polymerase to extend a template-bound oligonucleotide trigger and a nicking endonuclease to free the newly made product. The most common example of this reaction scheme is the exponential amplification reaction, or EXPAR^4^. Isothermal oligonucleotide amplification reactions are widely incorporated into DNA circuits and logic gates^2,5^, miRNA detection^6,7^, aptamer-based analyte detection^8^, RNA detection^9^, and genomic DNA detection^10^, and are thus broadly used across a variety of disciplines. While many works have shown the utility of oligonucleotide amplification, a base biphasic switch-like amplification reaction has not yet been reported.

Switch-like responses to input stimuli are ubiquitous in nature. This switching behavior is common in cell signaling, transcription, and genetic regulatory networks; it is commonly accepted that these switches react decisively to a true signal while filtering out noise^11^. Several studies have reported switch-like behavior in synthetic biochemical systems. Ion channels can be repurposed into biosensor switches by preventing channel dimerization in the presence of an target antigen, thus turning on in the presence of target^12^. DNA oscillators can switch between an “on” and “off” state by combining DNA degradation with a DNA amplification reaction^5^. It was noted that this oscillatory effect could be achieved through non-linear DNA amplification instead of non-linear DNA degradation, but the former was difficult to obtain and manipulate and was therefore not an option when creating a DNA circuit. Structure-switching sensors such as aptamers^13^ and molecular beacons^14,15^ change conformation in the presence of a specific target molecule. When properly designed, structure-switching biosensors can also create Hill-type ultrasensitive kinetics: biosensors with two cooperative binding sites produce an ultrasensitive response if the affinity of the target for the second site is altered by target association to the first site^16−19^. These exciting biomimetic systems typically produce outputs with nanomolar trigger inputs. A single cell can contain as few as 10 microRNA molecules per cell^20^, and clinically relevant DNA and RNA concentrations range from hundreds of picomolars to attomolar in range^21^. Clinically relevant protein concentrations are often in the femtomolar range^22^. While these previous studies explored sensors that are controlled switches, most do not have the subsequent high-gain amplification required for low target concentrations.

We present a rapid isothermal nucleic acid amplification method with an endogenous switching mechanism. The method exploits a naturally occurring stall in the amplification reaction, which produces a low-level signal. Upon surpassing a threshold, the reaction enters a high-gain second phase “burst”, producing an oligonucleotide concentration that ranges from ten to one hundred times the first phase plateau. Here we show the conditions required for entering the second phase as well as proposed reaction mechanisms driving two distinct types of switch-like reactions. Output kinetics can be tuned to control reaction acceleration in the second phase, resembling definitive switch turn-on. Additionally, reaction design using controlled DNA association thermodynamics give some control over first phase kinetics. Proteins^8,23−26^, genomic bacterial DNA^10^, viral DNA^27^, microRNA^28^, or mRNA^9^ can be transduced into many oligonucleotide triggers, making this technique applicable to a broad range of biological sensors.

## EXPERIMENTAL SECTION

### Reagents

UltraPure™ Tris-HCI pH 8.0, RNase free EDTA, RNase free MgCl_2_, RNase free KCl, Novex™ TBE Running Buffer (5X), 2X TBE-Urea Sample Buffer, Novex™ TBE-Urea Gels, 15%, SYBR^®^ Gold Nucleic Acid Gel Stain, and SYBR^®^ Green II RNA Gel Stain were purchased from Thermo Fisher Scientific (Waltham, MA). Nuclease-free water and oligo length standard 10/60 were purchased from Integrated DNA Technologies, Inc. (Coralville, IA). Nt.BstNBI nicking endonuclease, Bst 2.0 WarmStart^®^ DNA Polymerase, 10x ThermoPol I Buffer, dNTPs, BSA, and 100 mM MgSO_4_ were purchased from New England Biolabs (Beverly, MA).

Oligonucleotides were ordered from two different sources to avoid trigger contamination in templates. Desalted amplification templates were purchased from Integrated DNA Technologies (Coralville, IA) suspended in IDTE Buffer at a concentration of 100 μM. Templates were modified with an amino group on the 3’ end to prevent template extension. All desalted trigger oligonucleotides were purchased from Eurofins Genomics (Louisville, KY) suspended at a concentration of 50 μM in TE Buffer. Triggers were diluted in nuclease-free water in a separate room to prevent contamination.

### Template design and thermodynamics

Thermodynamics of the template stem loops were determined using the Mfold web server^29^, an open source software that uses empirical free energies of DNA hybridization^30^ that have been corrected for salt concentration^31^ (http://unafold.rna.albany.edu/?q=mfold). The free energies of association between the template and trigger, template and elongated trigger, product dimers, and double stranded templates were determined using the DINAmelt application, two-state melting (http://unafold.rna.albany.edu/?q=DINAMelt/Two-state-melting). To determine the free energy of toehold association, the software input was the sequence of the toehold and the toehold reverse compliment. All settings used were kept at the default software parameters, except for temperature (55°C) and salt concentration ([Na^+^] = 60mM, [Mg^++^] = 6mM). Templates were rejected if the structure with the lowest free energy was not the expected looped structure. Templates that are triggered by triggers ≤ 8 nucleotides were not included in the final analysis, as they gave little to no amplification under our reaction conditions (LS2 sp and LS2 st, Table SI 1).

### Biphasic Amplification Reactions

The amplification reaction mixture contained 1x ThermoPol I Buffer [20 mM Tris-HCl (pH 8.8), 10 mM (NH_4_)_2_SO_4_, 10 mM KCl, 2 mM MgSO_4_, 0.1% Triton^®^ X-100], 25 mM Tris-HCl (pH 8), 6 mM MgSO_4_, 50 mM KCl, 0.5 mM each dNTP, 0.1 mg/mL BSA, 0.2 U/μL Nt.BstNBI, and 0.0267 U/μL Bst 2.0 WarmStart^®^ DNA Polymerase. Bst 2.0 WarmStart^®^ DNA polymerase is inactive below 45°C; this decreases non-specific amplification before reaction initiation and theoretically increases experimental reproducibility. Templates were diluted in nuclease-free water and added at a final concentration of 100 nM. SYBR Green II (10,000x stock in DMSO) was added to the reaction mixture to a final concentration of 5x. Reactions were prepared at 4°C, and triggers and templates were handled in separate hoods to prevent contamination. Triggers were diluted in nuclease-free water and added to positive samples to a final concentration of 10 pM unless otherwise indicated; negative controls contained no trigger. For each experiment, two controls were prepared: a no-template control (NTC) sample containing no template, and a no-enzyme control sample containing no enzymes. Reactions were run in triplicate 20 μL volumes. Fluorescence readings were measured using a BioRad CFX Connect Thermocycler (Hercules, CA). Measurements were taken every 20 seconds with a 12 second imaging step. Reactions were run for either 150 or 300 cycles of 32 seconds at 55°C. The mixture was heated to 80°C for 20 minutes to deactivate enzymes, followed by 10°C for five minutes to cool the samples. Completed reactions were stored at −20°C for further analysis.

### Data analysis

Real-time reaction traces were analyzed with custom software using Matlab (Natick, MA). Details on calculation of inflection points and maximum reaction rates can be found in the SI (Figure SI 5, custom Matlab analysis software). The ratios between maximum reaction rates and inflection points were calculated from at least two experiments with three experimental replicates each. When appropriate, data from two experiments were averaged using a weighted average. Spearman’s rank-order correlations and p-values were determined using the function “corr” with the type selected as “Spearman” in Matlab (Natick, MA). Further details of statistical analysis can be found in the SI under “statistics”.

## RESULTS AND DISCUSSION

### Reaction pathways in the biphasic DNA amplification reaction

The biphasic DNA amplification reaction contains the same base components as the exponential amplification reaction for oligonucleotides (EXPAR)^4^. Both EXPAR and the biphasic DNA amplification reaction amplify a trigger sequence of ten to twenty base pairs in length at a single reaction temperature of 55°C through the action of a thermophilic polymerase and a nicking endonuclease. Both reactions non-specifically create product in the absence of initial DNA trigger, and exhibit similar separation between this non-specific amplification and specific trigger-initiated amplification under our tested conditions (Figure SI 6). The primary difference between the original EXPAR reaction and the biphasic oligonucleotide amplification reaction is the palindromic sequence within the DNA template that causes the template to fold into a looped configuration. The thermodynamics of the trigger binding and DNA association lie in a regime that creates a biphasic DNA amplification reaction; EXPAR-type DNA amplification using looped templates is found in literature^34−36^, but the biphasic kinetics have not yet been reported or analyzed.

Representative outputs of the oligonucleotide amplification reaction are shown in Figure 1. Despite the similarities in reaction components, the biphasic amplification reaction reported here is functionally distinct from all other EXPAR reactions. The first phase of the reaction resembles traditional EXPAR output, with an initial rise and a first plateau. After the first plateau, the biphasic reaction enters a high-gain second phase. This finding reveals that EXPAR can recover from the first plateau, a fact that was previously unknown. One template favored a linear configuration at the reaction temperature (LS3 lowpG2, T_m_ = 49.2°C) (Figure SI 2) and still gave biphasic output; this implied that while a palindromic region is necessary for biphasic output, a stable loop structure is not.

**FIGURE 1.**
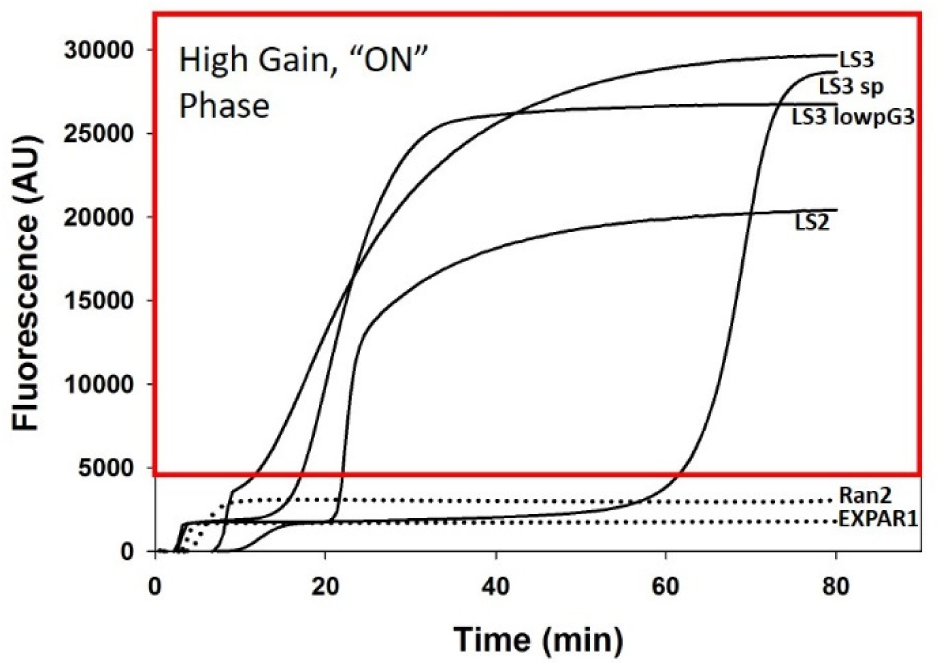
Representative biphasic amplification reaction output. DNA amplification output is correlated to fluorescence, which increases and plateaus at approximately the same level as previously reported optimized EXPAR reactions^32,33^ (dotted lines). Biphasic DNA amplification output is shown in solid lines. After a lag period, the DNA output jumps into a high gain “ON” region. Template DNA names are labeled next to corresponding output traces; template sequences can be found in the Table SI 1.

The mechanism behind the switch-like oligonucleotide amplification reaction is likely driven by multiple phenomena, as shown in Figure 2. The DNA template is composed of two copies of the complementary sequence joined by a ten-nucleotide nicking enzyme recognition site. The template contains a 3’ amine group to prevent extension of the template, a 3’ toehold, a palindromic sequence, the nickase recognition site, the repeated 5’ toehold, and the repeated palindromic sequence. The palindromic region causes the template to fold into a looped configuration (top panel, Figure 2). Triggers for these templates consist of the toehold complement and the template palindrome. When trigger binds to the 3’ end of the template and unwinds the template loop, the DNA polymerase extends the strand and the nicking enzyme recognition site is created. The nickase then nicks the growing strand. The polymerase can extend at this nick and will displace downstream trigger that has not dynamically dissociated from the template. Two distinct template types are discussed here: Type I templates with dynamic trigger:template association at the reaction temperature (Tm < 60°C), and Type II templates with stable trigger:template association at the reaction temperature (Tm > 60°C). When the displaced trigger freed it can prime other templates, leading to exponential amplification. The amplification therefore produces both triggers and long triggers that contain the nickase recognition site on their 3’ end.

**FIGURE 2.**
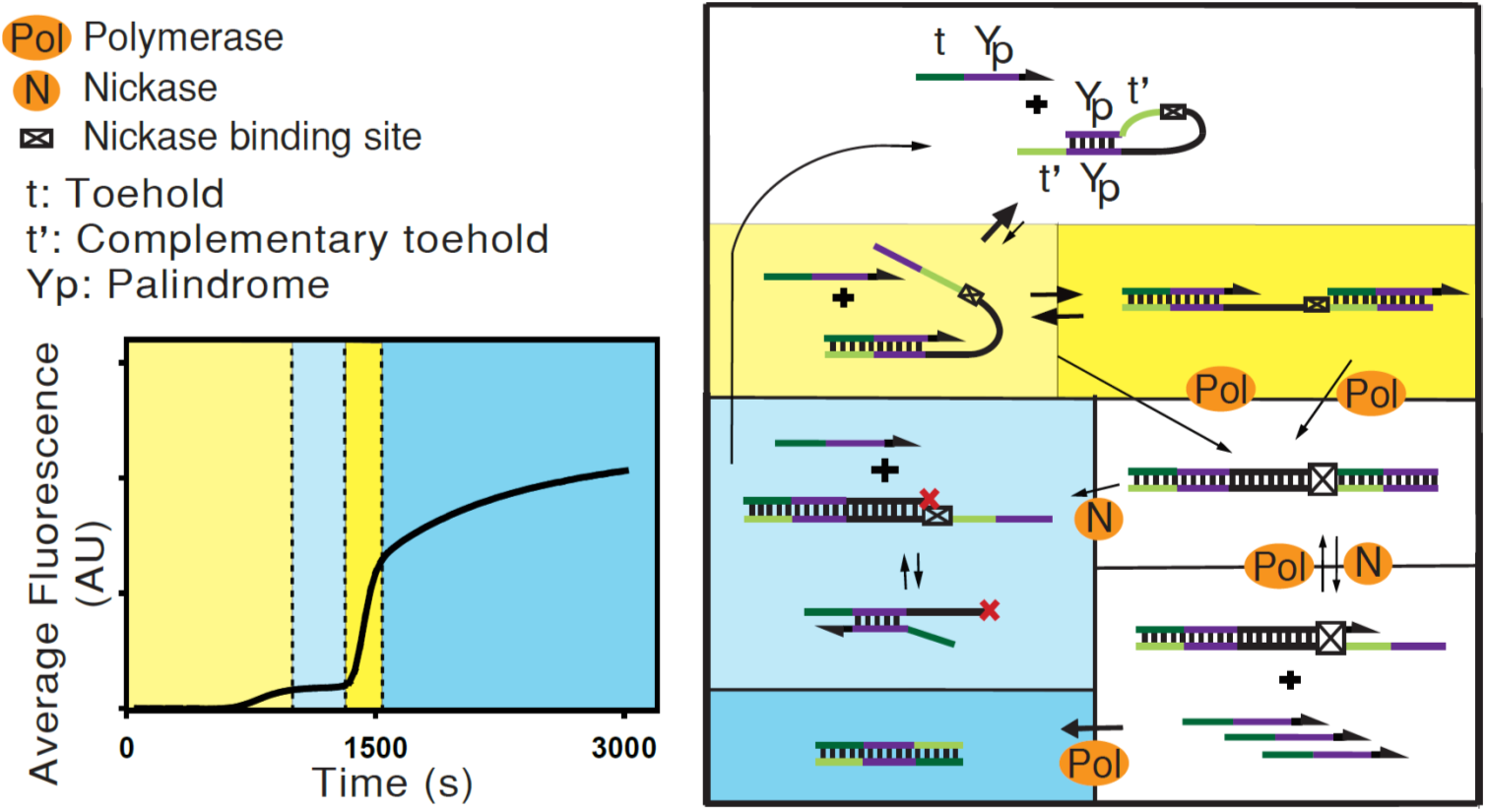
Biphasic DNA amplification reaction. The cartoon depicts potential reaction pathways in the biphasic DNA amplification reaction. The amplification requires a looped DNA template with two palindromic sequences (Yp), two toeholds (t’), and a restriction site (⊠), as well as polymerase and nickase enzymes. The reaction amplifies a DNA trigger with a reverse compliment to the template toehold (t) and the palindromic region (Yp). Arrows show extendable 3’ ends of the DNA; the 3’ end of the template is blocked with an amine group to prevent non-specific elongation. The trigger can bind to either toehold region t’ and strand displace the palindromic region Yp, thus opening the loop (yellow). A polymerase can then extend the trigger and create the recognition site for a nicking endonuclease, as well as an identical trigger. The nickase then cuts the top strand, freeing the newly created trigger to bind other templates. The loop can also remove the long trigger and close with the aid of triggers, which can bind the long trigger and facilitate loop closure. This may be vital to remove “poisoned” long triggers that cannot amplify and block further trigger amplification on the template (light blue). The palindromic region can also cause trigger dimerization, after which the toehold regions can be filled by the polymerase (dark blue); this removes trigger molecules from further amplification cycles. Colored regions on the amplification curve correlate with the proposed dominant reaction mechanism in each reaction phase.

The presence of the palindromic sequence on the triggers and template produces several new reaction pathways; the proposed dominant reaction pathways in each phase of the reaction are denoted by color in Figure 2. Long triggers contain the trigger sequence and the newly elongated nickase recognition site. The trigger can catalyze removal of this long trigger through association of the palindromic region of the long trigger and the trigger (Figure 2, light blue), and subsequent loop closure. The association of the template and the first trigger molecule will open the loop, which both aids and stabilizes a second trigger association. For most templates, the looped configuration is more stable than the open, trigger-bound configuration (Table SI 2). The loop structure of the templates with two toehold regions may possibly create cooperative binding between the triggers and the looped template: the association of the first trigger will open the loop and make the second trigger association more thermodynamically favorable. The palindromic section of the triggers can also associate, be extended by the polymerase, and create inert triggers unable to further replicate; this pathway has been previously discussed^34^ (Figure 2, dark blue). We hypothesize that these new reaction pathways create the unique features of our amplification reaction.

### Properties of the first reaction phase

The first reaction phase resembles the base EXPAR reaction, with a low-gain reaction phase followed by a plateau. Inflection points are traditionally used as a surrogate for EXPAR reaction kinetics, so the kinetics of the first phase of the biphasic reaction were given by the first inflection point. Thermodynamics of the looped DNA template and 3’ toehold association are correlated with the first-phase reaction kinetics of Type I templates: templates with a 3’ toehold free energy that is lower than the loop free energy have rapid first phase kinetics (193±17 seconds) and templates with a 3’ toehold free energy that is higher than the loop free energy have slower first phase kinetics (780±90 seconds). Type II templates have different behavior and showed only modest correlation between first-phase reaction kinetics and DNA association thermodynamics (Figure SI 1).

While the reaction plateau stall was previously attributed to loss of nickase integrity^4^, the recovery of the reaction after the first plateau when using palindromic templates suggests otherwise. Recently others have hypothesized that templates could be “poisoned” due to polymerase errors that render the DNA strand bound to the template unextendable^37^. We hypothesize that poisoned templates could cause the plateau seen in the original EXPAR reaction and the first plateau in the biphasic amplification reaction. We estimated the plateau trigger concentration to be between 1-10 μM, with the plateau concentration of a standard EXPAR reaction being 5.58μM (Table SI 3), and the plateau concentration of a selected looped template LS3 at 2.5μM (Figure SI 3). The plateau concentration of trigger molecule is therefore between 10 and 100 times greater than the template concentration of 0.1μM. Due to the rapid template inactivation after at most 100 cycles of extension and nicking it is unlikely that polymerase error causes this plateau, given error rates of polymerases such as *Bst* DNA polymerase that lack the 3′ → 5′ exonuclease domain are approximately 10^−4^ ^38^. We hypothesize that the plateau is due to noncanonical behavior of the nickase enzyme that leaves a long unextendable trigger, as the nickase is operating in suboptimal conditions when compared to the polymerase^33^. It is also possible that a fully elongated trigger poisons the template; however, the mechanism behind the template poisoning is beyond the scope of this study.

### Properties of the second reaction phase

After the first plateau, the amplification enters a high-gain second phase followed by a second plateau. The amplification does not exit the first plateau unless there is a palindromic region in the template; we hypothesize that template rescue is aided by trigger association to the long “poisoned” triggers in conjunction with subsequent loop closure. This trigger-dependent rescue would prevent the long trigger from reassociating with the template, particularly after polymerase extension of the 3’ trigger end. The trigger could also bind the template and prevent reassociation of the long trigger. These events would aid in the loop closure and template rescue (Figure 2, light blue panel). After exiting the first plateau many of the templates exhibit rapid second phase kinetics, marked by a large jump in reaction product that can exceed first phase reaction kinetics. It is possible that second phase could be driven by homotropic allosteric cooperativity^17^; the trigger can bind either toehold as seen in Figure 2. The template loop structure is stable when compared to the trigger:template association (Table SI 2), and the accumulation of reaction products may possibly shift templates to an open, amplification competent state and produce nonlinear reaction kinetics.

The subsequent second plateau is caused by exhaustion of reaction components and a buildup of inhibitory reaction products, as shown in the dark blue panel of Figure 2. This effect of inhibitory products was previously described when using EXPAR reactions and a palindromic looped template^34^. The final output of the second phase is approximately the size of the DNA triggers and elongated triggers as seen in PAGE analysis of reaction products; for more details see the Supplementary Information (Figure SI 4). This rescue of the poisoned templates allows the reaction to produce 10-100 times more endpoint reaction product as measured by calibrated SYBR II fluorescence. Endpoint product concentration ranged from 7.8 - 116.9 μM, with several reaction products exceeding 100 μM during the second plateau (Table SI 3).

### Reaction response to initial trigger concentration

We measured reaction output in real-time with varying initial trigger concentrations using the ssDNA binding dye SYBR II for fluorescent readout. Figure 3A, B, and C show representative real-time fluorescent traces. The average background fluorescence from samples with original trigger concentrations ≤10nM was subtracted from all samples. It is important to note that absolute fluorescence units are arbitrary; indeed, the resolution between 1μM and 10μM fluorescence levels is smaller than the background noise between samples in Figure 3A. This fluorescence variation does not affect the shape and inflection points of the graph, however. Traces are shown for three different template types: a traditional EXPAR linear template (EXPAR1^32^), a type I template (LS2 lowtG), and a type II template (LS3), respectively. The traditional EXPAR template was chosen from 384 published sequences as it had the highest separation between positive and negative controls. Fits were performed on inflection points for initial trigger concentrations between the lowest detectable concentration (100fM or 1pM) and the highest concentration below the first plateau (1μM).

**FIGURE 3.**
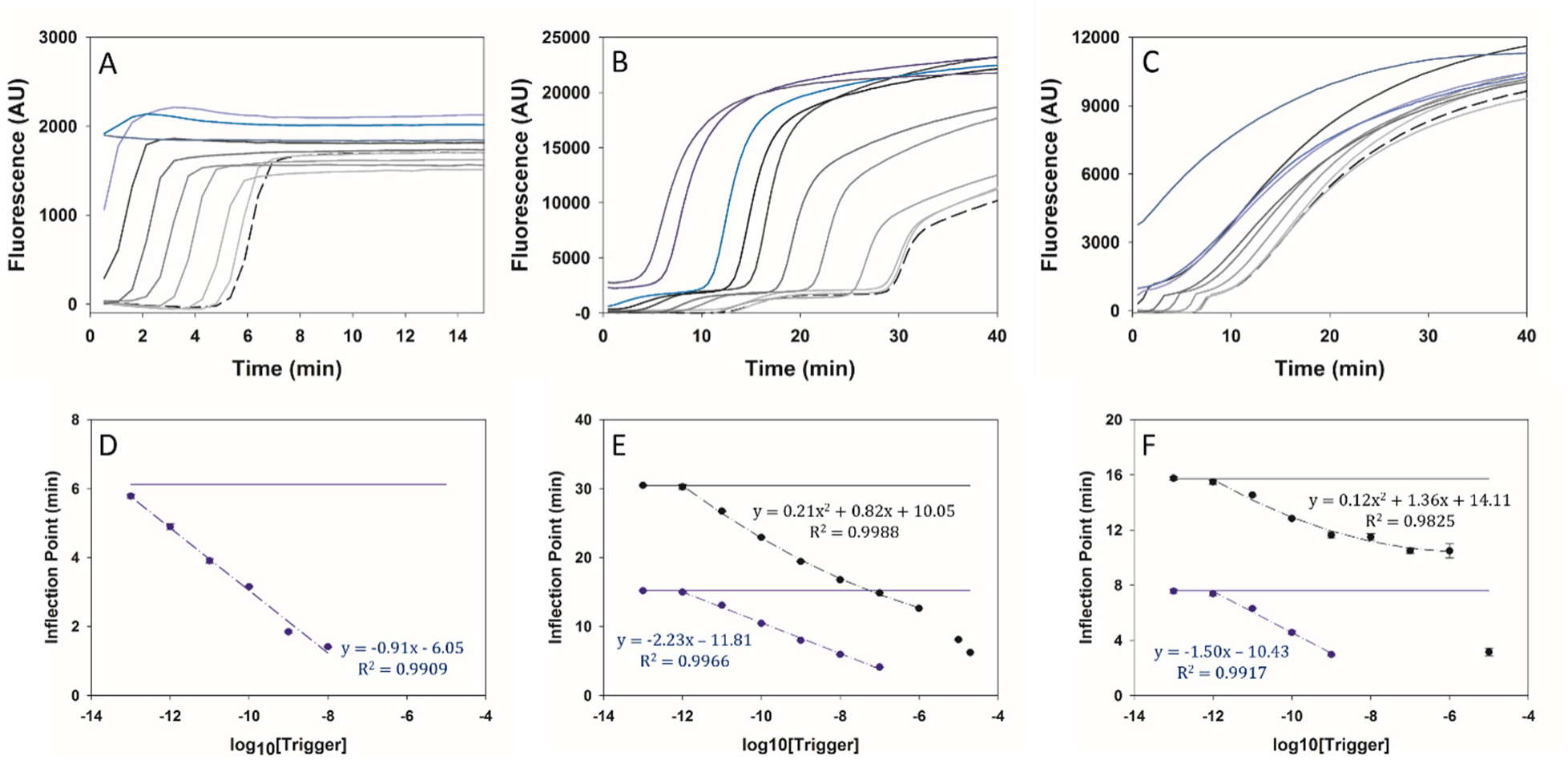
Correlation between amplification initiation and trigger oligonucleotide concentration. The real-time reaction output for three representative templates shows the dependence of the reaction on initial trigger concentration, with fluorescence correlated to the produced DNA trigger. Initial trigger concentrations were increased tenfold between 100 fM and 10 μM unless otherwise indicated; blue traces did not have measurable first reaction phases and darker color indicates higher initial trigger concentrations. A) Dilution series of a standard EXPAR template (EXPAR1^32^) do not enter the second phase, even at high concentrations of trigger (10 μM) B) Dilution series of the representative Type I template LS2 lowtG, which includes an extra trace at 20μM initial trigger concentration. C) Dilution series of the representative Type II template LS3. Calculated inflection points are shown for D) EXPAR1, E) LS2 lowtG, and F) LS3. Dashed lines show fits; blue symbols show first inflection points and black symbols show the second inflection point. Error bars represent standard deviation of experimental triplicates.

The calculated first and second inflection points are shown in Figure 3 D-F. The traditional EXPAR template (Figure 3 A,D) did not enter the second phase, even when initial trigger concentration (10μM) was above the plateau concentration of this template (5.58μM, Table SI 3). Inflection points of the traditional template were linearly correlated with the log_10_ of the original trigger concentration as expected^33^. Inflection points in the first phase for both the type I and II looped templates were also linearly correlated with the log_10_ of the original trigger concentration, but the correlation appeared slightly non-linear during the second phase (Figure 3 E,F). When the initial concentration of trigger exceeded the concentration at the first plateau, the inflection points appear to occur earlier than the fit lines predicted, which is expected for a hill-type reaction. The type I template initiated in the first plateau for initial trigger concentrations >1 μM and showed a short lag in amplification that was not present in the type II templates. The type I template also has a higher second plateau fluorescence level for greater initial concentrations of trigger, although it is unclear why this occurred. The type II template initiated in the second phase when the initial trigger concentration was 10 μM, which was above the measured plateau concentration (2.5 μM, Figure SI 3). This demonstrated the trigger dependence of the plateau and suggested that entering the second reaction phase was dependent on trigger concentration.

As with traditional EXPAR, the limit of detection for the DNA trigger was determined by the nonspecific amplification rates. The optimized traditional template was kinetically distinct from the negative control at 100fM of initial trigger, and the looped templates were kinetically distinct from the negative control at approximately 1pM of initial trigger. Nearly all looped templates tested could distinguish between 0 and 10 pM initial trigger concentrations (Figure SI 6).

### Weakening loop thermodynamics within templates

We expect that weakening the loop structure would accelerate reaction kinetics in the first phase; a weaker loop will open and amplify faster than a strong loop. To examine this phenomenon, the free energy of the looped template structure was altered by adding long random sequences of 4-8 nucleotides into the loop before the nickase recognition site. This modification held the palindrome, toehold, and trigger sequences constant while weakening the template loop structure. The only additional thermodynamic parameter this modified was an increase in stability of the long trigger:template complex, which forms after the initial elongation of a template-bound trigger. Long triggers included the original trigger sequence, the nickase recognition site, and the additional long random sequences. The first inflection points of these new long random sequence (lrs) templates were divided by the first inflection point of the base template with no lrs; a relative first inflection point of one denotes a template that was not affected by the weakened loop.

These weakened template loops produced surprising and revealing results. For type I template LS2, decreasing the strength of the loop appeared to cause the loop to open faster, although the p values from a Holm-Bonferroni t-test were not significant (p<0.09 and p<0.06, without correction). Type I template LS2 lowtG had similar or slightly slower trigger production in the first phase when the loop was weakened (Figure 4, blue bars). Surprisingly, decreasing the strength of the type II template loops slowed the first reaction phase for every template tested, with significant reaction delay for LS3 lt-1 templates and LS3 lrs-4 (Figure 4, grey bars). We hypothesize that the increased stability of the long triggers caused this phenomenon; these long triggers were more stable and therefore more difficult to remove. This observed decrease in the first phase reaction rate supported the hypothesis that templates were deactivated by unextendable “poisoned” complimentary strands, and that type II templates were more susceptible to this phenomenon than type I templates.

**FIGURE 4:**
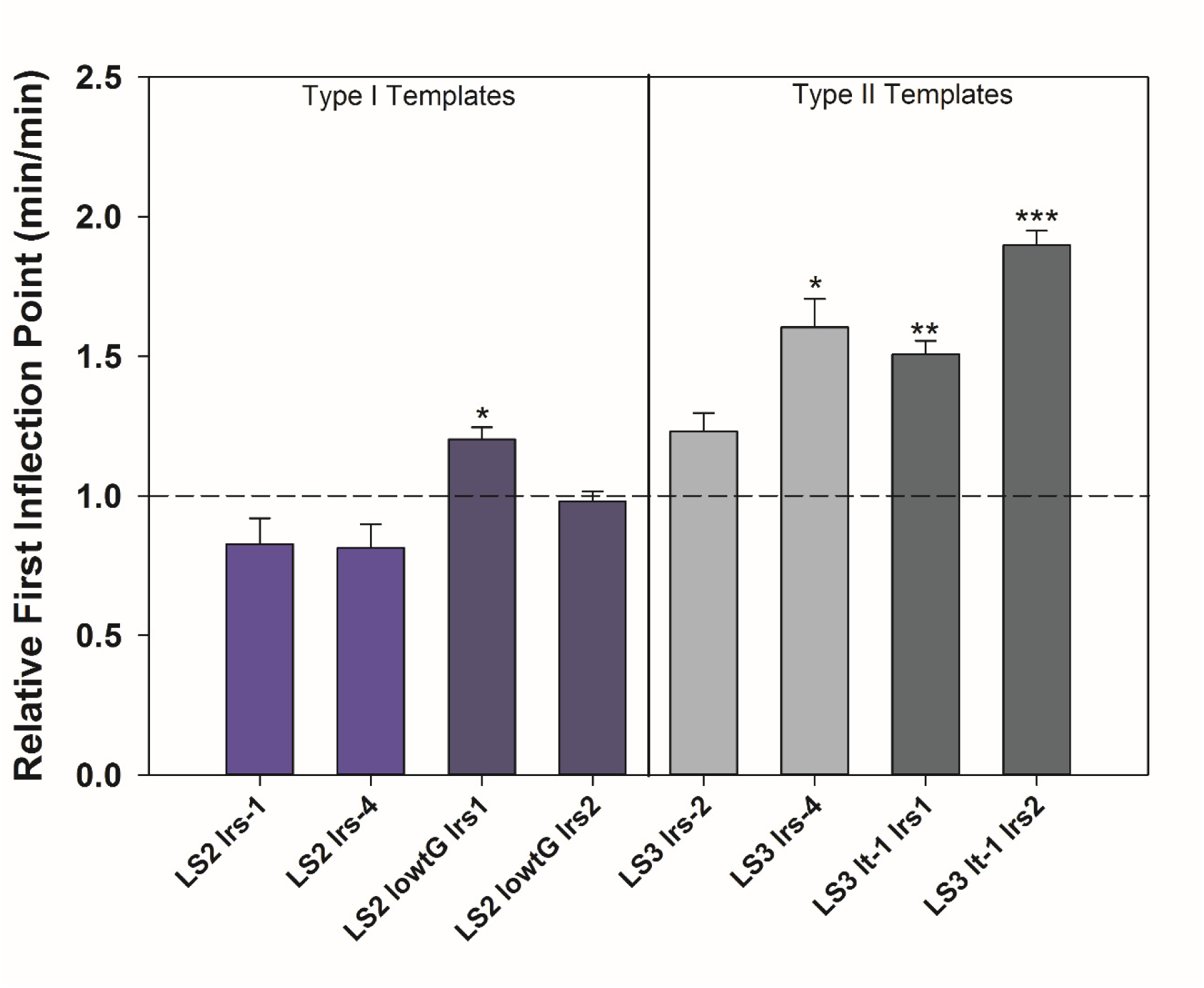
Weakening the loop structure of templates can result in slower reaction kinetics. Long random sequences (lrs) were added to four base looped templates after the nickase recognition site, resulting in a weaker template loop that produces the same product. The relative first inflection point is the average first inflection point divided by the average first inflection point of the base template without lrs; a value greater than one therefore signifies reduced first phase reaction kinetics. While a weakened loop has a modest effect on Type I templates (trigger Tm<60°C), Type II templates (trigger Tm>60°C) with weakened loops are much slower than their base template. Error bars represent standard deviations from at least three independent experiments, which all contained experimental replicates. * p<0.05, **p<0.01, ***p<0.001, Holm - Bonferroni t-test.

### Analyzing acceleration in the second reaction phase

Rapid acceleration in the second phase would be beneficial if these reactions are used as a digital readout, because a large jump in the second phase resembles definitive switch turn-on. We further analyzed DNA amplification kinetics for their ability to accelerate in the second phase, and to determine if relative second phase kinetics correlated with DNA association thermodynamics. The second phase acceleration was defined as the ratio of the maximum reaction rate in the second phase to the maximum reaction rate in the first phase; Figure SI 5 gives details of this calculation. We hypothesized that cooperative binding of the trigger to the two toehold binding sites could possibly cause a rapid increase in reaction kinetics in the second phase. Hill coefficients of a homotropic cooperative receptor increase with the ratio between dissociation constants of the first and second binding events; a large difference in stability between the first and second ligand associations will result in a larger Hill coefficient and greater Hill behaviour^17^. This is qualitatively intuitive: the more relative stability that the first association provides, the greater the benefit from having a higher concentration of ligand. In our system, this corresponds to the amplification incompetent state (a closed template) moving to an amplification competent state (an open template) through dual trigger binding. We characterized the relative dissociation of the first and second trigger binding events by the difference in the free energy of the first binding event Δ*G*_5′*toehold*_ + Δ*G*_3′*toehold*_ + Δ*G*_*palindrome*_ - Δ*G*_*loop*_ and the second binding event Δ*G*_*trigger*:*template*_. A larger value signified a greater difference between the stability of the first and second trigger associations to the template.

The two template types showed distinct behavior. Type I templates showed significant correlation between the difference between the first and second trigger binding event and the reaction acceleration in the second phase (Spearman’s Rho = 0.9667, p<1.7x10^−4^), while Type II templates did not (Spearman’s Rho = 0.6437, p<0.10) (Figure 5A). The association of one trigger with these templates was thermodynamically unfavorable when compared to the stable loop structure (Table SI 2), but upon association the trigger will open the loop structure and switch the receptor to a binding competent state (Figure 2A, panel 1).

**Figure 5:**
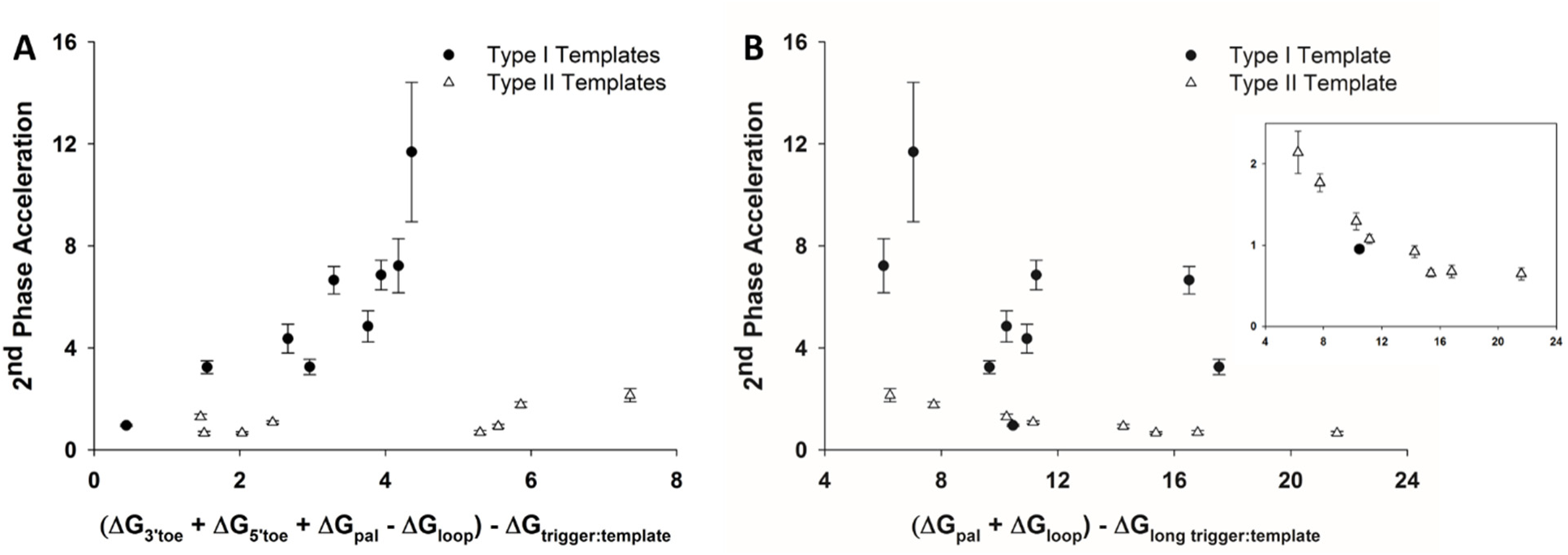
Acceleration of trigger production in the second phase vs. DNA association thermodynamics. Type I and type II templates show two distinct behaviors in the second phase. A) Type I templates have triggers that can dynamically dissociate from the template at the reaction temperature (Tm > 60°C). If the cooperativity of trigger binding to the two open toeholds contributed to reaction acceleration in the second phase, then the thermodynamic difference between the first and second trigger binding events (Δ*G*_5′*toehold*_ + Δ*G*_3′*toehold*_ + Δ*G*_*palindrome*_ -Δ*G*_*loop*_) - Δ*G*_*trigger*:*template*_ would correlate with accelerating kinetics in the second phase. This is true of type I templates (Spearman’s Rho = 0.9667, p<1.7x10^−4^), but not type II templates (Spearman’s Rho = 0.6437, p<0.10). B) Type II templates have triggers that are stable at the reaction temperature (Tm > 60°C), making loop closure and long trigger removal more difficult. Long trigger removal, as described in Figure 2, is approximated by Δ*G*_*long trigger*:*trigger*_ + Δ*G*_*loop*_ - Δ*G*_*long trigger*:*template*_. This correlates with second phase acceleration of type II templates (Spearman’s Rho = −0.9762, p<4.0x10^−4^), but not type I templates (Spearman’s Rho = −0.3333, p<0.39). Inset is rescaled to show a zoomed in graph of type II template second phase acceleration.

Type II templates appear to have different dominant reaction pathways. Long trigger removal was hypothesized to be driven by loop closure, which would be hindered by the presence of a stable triggers that remained on type II templates after nicking. The low acceleration in the second phase seen in type II templates could possibly be due to hindered removal of long poisoned triggers. Figure 5B supports this hypothesis: the parameter Δ*G*_*long trigger*:*trigger*_ + Δ*G*_*loop*_ - Δ*G*_*long trigger*:*template*_ approximates thermodynamics of long trigger removal through trigger association to the long trigger and subsequent loop closure. A larger value corresponds to more stable long triggers. Type II templates significantly correlate with this thermodynamic parameter, with the stability of the long triggers inversely proportional to the reaction acceleration in the second phase (Spearman’s Rho = −0.9762, p<4.0x10^−4^). This correlation was not significant when analyzing type I templates (Spearman’s Rho = −0.3333, p<0.39). These observations support the concept of two distinct templates types.

## CONCLUSIONS

We report a novel, biphasic DNA amplification reaction with a low gain first phase followed by a high gain second phase. The first phase resembles the popular oligonucleotide amplification reaction EXPAR, which operates on similar principles but uses a linear DNA template without a palindromic sequence. We hypothesize that the accumulation of “poisoned” templates with bound unextendable long triggers may slow the reaction and cause the first plateau seen in most palindromic templates and all EXPAR templates. The presence of a palindrome in the template appears to rescue the reaction from this plateau, even when the loop structure is not stable at the reaction temperature (Figure SI 2). While palindromes can rescue the reaction from the first plateau, not all palindromic templates showed this first plateau (Figure SI 7). Several highly stable loops had slow kinetics and unmeasurable plateaus, and it was unclear if they entered the second phase. Two templates with vanishingly small plateaus that entered the second phase both had relatively stable trigger:template complexes, although with only two templates it is not clear from this data the exact parameters that cause the plateau phase to effectively disappear.

The reactions investigated here fall into two distinct categories: type I templates had triggers that will dynamically dissociate from the template after nicking (Tm<60°C), while type II templates had stable triggers that were more likely to remain until strand displacement by the polymerase (Tm>60°C). DNA association thermodynamics related to the first phase reaction kinetics and second phase acceleration within the two template types, but did not show the same correlations between template types. Long template-bound triggers appeared to slow the reaction, particularly for type II templates which had stable triggers bound after nicking. We hypothesize that these effects caused a smaller acceleration in the second phase for type II templates, which typically did not show the same switch-like jump in second phase product concentration when compared to type I templates. These observations suggested that the two types of templates had different dominant reaction pathways, and provided important design considerations to tune the reaction output during each phase.

These biphasic reactions require further investigation and optimization. Mathematical modeling of reaction kinetics and a full mechanistic understanding of the reaction phases would aid reaction design. As with many isothermal amplifications such as EXPAR, this reaction also produces non-specific amplification occurring at long reaction times in the absence of an oligonucleotide trigger. Non-specific amplification will increase the limit of detection and can decrease the experimental robustness. Optimizing solution conditions, adding ssDNA binding proteins and carbon sheets^39^, or adding other small molecules^40^ can decrease non-specific amplification in EXPAR reactions, and would likely also be applicable in this reaction. Degradation of the reaction product was previously used to create a bistable switch from an EXPAR-type reaction^5^, and could be extended to suppress non-specific amplification or create threshold-based detection for targets above a chosen concentration. Inhibition or degradation of reaction products may also create a true bistable switch that could repeatedly turn “off” and “on”.

We have demonstrated a novel new oligonucleotide amplification reaction with a two-stage output that is dependent on the released oligonucleotide trigger molecule. This biphasic DNA amplification reaction is a simple, one-step isothermal amplification reaction; reactions of this type have gained popularity as they do not require temperature cycling and therefore require less energy, hardware, and time^1,41^. We have described a thermodynamically-based reaction design framework to approximate first phase output, as well as to tune the reaction acceleration in the switch-like second reaction phase. The reaction can report on a variety of analytes: specific proteins^8,23−26^, genomic bacterial DNA^10^, viral DNA^27^, microRNA^28^, or mRNA^9^ can continuously create input trigger oligonucleotides, making the biphasic DNA amplification reaction broadly applicable to a variety of target molecules. When combined with single molecule amplification, this technique has the potential to be quantitative through digital amplification and detection^42^. The biphasic nature of this reaction makes it well suited for recognition of low-concentration molecules in biological samples, DNA logic gates, and other molecular recognition systems.

## Acknowledgements

This publication was supported by the National Center for Advancing Translational Sciences of the National Institutes of Health [UL1 TR002319] and the Montana Research and Economic Development Seed Funds.

## References

(1) McCalla, S. E.; Tripathi, A. Annual review of biomedical engineering 2011, 13, 321–343.

(2) Zhang, D. Y.; Seelig, G. Nature chemistry 2011, 3, 103–113.

(3) Zhang, D. Y.; Winfree, E. Journal of the American Chemical Society 2009, 131, 17303–17314.

(4) Van Ness, J.; Van Ness, L. K.; Galas, D. J. Proceedings of the National Academy of Sciences 2003, 100, 4504–4509.

(5) Montagne, K.; Gines, G.; Fujii, T.; Rondelez, Y. Nature Communications 2016, 7, 13474.

(6) Chen, S. X.; Seelig, G. Journal of the American Chemical Society 2016, 138, 5076–5086.

(7) Duan, R.; Zuo, X.; Wang, S.; Quan, X.; Chen, D.; Chen, Z.; Jiang, L.; Fan, C.; Xia, F. Journal of the American Chemical Society 2013, 135, 4604–4607.

(8) Zhang, Z.-z.; Zhang, C.-y. Analytical Chemistry 2012, 84, 1623–1629.

(9) Zhao, Y.; Zhou, L.; Tang, Z. Nat Commun 2013, 4, 1493.

(10) Roskos, K.; Hickerson, A. I.; Lu, H.-W.; Ferguson, T. M.; Shinde, D. N.; Klaue, Y.; Niemz, A. PLoS ONE 2013, 8, e69355.

(11) Ferrell, J. E. Trends in biochemical sciences 1996, 21, 460–466.

(12) Cornell, B. A.; Braach-Maksvytis, V.; King, L.; Osman, P. Nature 1997, 387, 580.

(13) Zuo, X.; Song, S.; Zhang, J.; Pan, D.; Wang, L.; Fan, C. Journal of the American Chemical Society 2007, 129, 1042–1043.

(14) Vallée-Bélisle, A.; Ricci, F.; Plaxco, K. W. Proceedings of the National Academy of Sciences 2009, 106, 13802–13807.

(15) Tyagi, S.; Kramer, F. R. Nature biotechnology 1996, 303–308.

(16) Simon, A. J.; Vallée-Bélisle, A.; Ricci, F.; Plaxco, K. W. Proceedings of the National Academy of Sciences 2014, 111, 15048–15053.

(17) Simon, A. J.; Vallée-Bélisle, A.; Ricci, F.; Watkins, H. M.; Plaxco, K. W. Angewandte Chemie 2014, 126, 9625–9629.

(18) Ricci, F.; Vallée-Bélisle, A.; Simon, A. J.; Porchetta, A.; Plaxco, K. W. Accounts of Chemical Research 2016, 49, 1884–1892.

(19) Rossetti, M.; Porchetta, A. Analytica Chimica Acta 2018.

(20) Zhou, Y.; Huang, Q.; Gao, J.; Lu, J.; Shen, X.; Fan, C. Nucleic acids research 2010, 38, e156–e156.

(21) Kelley, S. O. ACS Sensors 2017, 2, 193–197.

(22) Rissin, D. M.; Kan, C. W.; Campbell, T. G.; Howes, S. C.; Fournier, D. R.; Song, L.; Piech, T.; Patel, P. P.; Chang, L.; Rivnak, A. J. Nature biotechnology 2010, 28, 595–599.

(23) Tang, W.; Zhang, T.; Li, Q.; Wang, H.; Wang, H.; Li, Z. RSC Advances 2016, 6, 89888–89894.

(24) Qiu, T.; Wang, Y.; Yu, J.; Liu, S.; Wang, H.; Guo, Y.; Huang, J. RSC Advances 2016, 6, 62031–62037.

(25) Nutiu, R.; Li, Y. Journal of the American Chemical Society 2003, 125, 4771–4778.

(26) Wieland, M.; Benz, A.; Haar, J.; Halder, K.; Hartig, J. S. Chemical Communications 2010, 46, 1866–1868.

(27) Tan, E.; Erwin, B.; Dames, S.; Voelkerding, K.; Niemz, A. Clinical chemistry 2007, 53, 2017–2020.

(28) Zhang, X.; Liu, C.; Sun, L.; Duan, X.; Li, Z. Chemical Science 2015, 6, 6213–6218.

(29) Zuker, M. Nucleic acids research 2003, 31, 3406–3415.

(30) SantaLucia, J. Proceedings of the National Academy of Sciences 1998, 95, 1460–1465.

(31) Peyret, N. Prediction of nucleic acid hybridization: parameters and algorithms; Wayne State University Detroit, 2000.

(32) Qian, J.; Ferguson, T. M.; Shinde, D. N.; Ramírez-Borrero, A. J.; Hintze, A.; Adami, C.; Niemz, A. Nucleic acids research 2012, 40, e87–e87.

(33) Tan, E.; Erwin, B.; Dames, S.; Ferguson, T.; Buechel, M.; Irvine, B.; Voelkerding, K.; Niemz, A. Biochemistry 2008, 47, 9987–9999.

(34) Fujii, T.; Rondelez, Y. ACS nano 2012, 7, 27–34.

(35) Padirac, A.; Fujii, T.; Estévez-Torres, A.; Rondelez, Y. Journal of the American Chemical Society 2013, 135, 14586–14592.

(36) Zambrano, A.; Zadorin, A. S.; Rondelez, Y.; Estévez-Torres, A.; Galas, J. C. The Journal of Physical Chemistry B 2015, 119, 5349–5355.

(37) Baccouche, A.; Montagne, K.; Padirac, A.; Fujii, T.; Rondelez, Y. Methods 2014, 67, 234–249.

(38) Lage, J. M.; Leamon, J. H.; Pejovic, T.; Hamann, S.; Lacey, M.; Dillon, D.; Segraves, R.; Vossbrinck, B.; González, A.; Pinkel, D. Genome Research 2003, 13, 294–307.

(39) Wang, J.; Zou, B.; Rui, J.; Song, Q.; Kajiyama, T.; Kambara, H.; Zhou, G. Microchimica Acta 2015, 182, 1095–1101.

(40) Mok, E.; Wee, E.; Wang, Y.; Trau, M. Scientific reports 2016, 6.

(41) Li, J.; Macdonald, J. Biosensors and Bioelectronics 2015, 64, 196–211.

(42) Zhang, K.; Kang, D.-K.; Ali, M. M.; Liu, L.; Labanieh, L.; Lu, M.; Riazifar, H.; Nguyen, T. N.; Zell, J. A.; Digman, M. A.; Gratton, E.; Li, J.; Zhao, W. Lab on a Chip 2015, 15, 4217–4226.

